# Insulin Resistance Mediates Crosstalk between Skeletal Muscle and the Heart to Prevent Cardiac Dysfunction

**DOI:** 10.1101/2020.11.14.383109

**Authors:** Dandan Jia, Jun Zhang, Xueling Liu, John-Paul Anderson, Zhenjun Tian, Jia Nie, Yuguang Shi

## Abstract

Obesity and type 2 diabetes milieus (T2DM) are the leading causes of cardiovascular morbidity and mortality. Although insulin resistance is believed to underlie these disorders, a paradox exists that obese patients with cardiovascular diseases have better prognoses relative to leaner patients with the same diagnoses, whereas improvement of insulin sensitivity by thiazolidines significantly increases the risk of heart failure in T2DM patients. Using mice with skeletal muscle-specific deletion of the insulin receptor gene (MIRKO), we resolved this dilemma by demonstrating that insulin resistance in the skeletal muscle prevents myocardial hypertrophy and dysfunction in response to diet-induced obesity. In spite of aggregating obesity, insulin resistance selectively protected the heart, but not other metabolic tissues, from mitochondrial dysfunction and insulin resistance in MIRKO mice, leading to significant attenuation of inflammation and apoptosis of cardiomyocytes. Together, our findings revealed an unexpected role of insulin resistance as a double edge sword, calling for reevaluation of ongoing efforts for targeting insulin resistance for the treatment of metabolic diseases.

**Significance statement:** Insulin resistance is commonly recognized as one of the major causes of T2DM and other aging-related chronic diseases. However, cumulative efforts in targeting insulin resistance for the treatment of T2DM and other metabolic disorders have not met with great success due to unwanted side effects on heart failure. Here we report that insulin resistance in skeletal muscle is a double edge sword. On one hand, insulin resistance in skeletal muscle aggregates obesity and its related metabolic syndromes, and on the other it protects the heart from the development of myocardial hypertrophy, dysfunction, and apoptosis in response to DIO. Our findings call for reevaluation of the ongoing strategies in targeting insulin resistance as a novel treatment for T2DM and other metabolic diseases.

## Introduction

Obesity causes insulin resistance which is implicated in the pathogenesis of metabolic diseases, including T2DM and cardiovascular diseases (CVD) (1). However, despite the success of many commonly used antihyperglycemic therapies to control hyperglycemia in T2DM, the prevalence of heart failure remains very high, raising the possibility that additional factors beyond blood glucose might contribute to the increased heart failure risk (2). Moreover, a paradox exists about the contradicting role of insulin resistance in the development of CVD. As the largest insulin sensitive organ in the body, skeletal muscle accounts for up to 85% of glucose disposal following a glucose infusion, yet its role in diabetes and CVD is far from clear (3). Circumstantial evidence suggests that insulin resistance in skeletal muscle might protect the heart in response to metabolic stress. In support of this notion, overweight and obese patients with CVD, including coronary heart disease and heart failure, have better short- and medium-term prognoses compared with leaner patients with the same cardiovascular diagnoses, as demonstrated by numerous studies and meta-analyses from across the world (4–6). Since most obese patients have metabolic syndrome including insulin resistance, the findings suggest that insulin resistance may benefit patients with CVD. Additionally, treatment of T2DM with thiazolidines (TZDs), an insulin sensitizer drug that was widely used for the treatment of T2DM, led to increased risk of heart failure, as evidenced by multiple meta-analysis of clinical trials (7) (8–10). This notion is further supported by experimental evidence from genetic studies of mouse models of insulin resistance and CVD. For example, insulin resistance in liver and islet β-cell led to severe hyperinsulinemia or T2DM, as demonstrated by the phenotypes of liver and islet β-cell specific insulin receptor gene knockout mice (11, 12). In contrast, MIRKO mice exhibited a relatively benign phenotype in glucose homeostasis (3). Moreover, myocardial specific deletion of the *OPA1* gene, which encodes a GTPase required for the fusion of the mitochondrial inner membrane, led to dilated cardiomyopathy and heart failure (13). This lethal phenotype was mitigated by either targeted deletion of *OPA1* in skeletal muscle or diet-induced obesity (DIO) (13). Together, these findings suggest that insulin resistance in skeletal muscle may benefit the heart under metabolic stress (13).

In this study, we tested the hypothesis that skeletal muscle insulin resistance protects the heart under metabolic stress. Using MIRKO mice as a mouse model of severe insulin resistance, we revealed an unexpected role of insulin resistance as the key mediator between skeletal muscle and the heart in response to metabolic stress. We showed that insulin resistance in skeletal muscle protected the heart from the development of cardiac hypertrophy and left ventricle dysfunction in response to DIO. We further showed that skeletal muscle insulin resistance also prevented mitochondrial dysfunction in response to DIO, leading to a significant improvement of insulin sensitivity and fatty acid oxidation in the heart, but not in other metabolic tissues. Our findings challenge long held concepts about the control of insulin resistance as the most detrimental factor in chronic diseases.

## Results

### MIRKO aggravates DIO-induced metabolic disorders, leading to hyperinsulinemia, glucose and insulin intolerance

Using the Cre-loxp system, we generated MIRKO mice with skeletal muscle specific deletion of the insulin receptor (IR) gene as previously reported (3). The approach led to a near total depletion of IR expression in skeletal muscle, as demonstrated by immunoblot analysis of insulin receptor α (IR-α) and β (IR-β) (Figure 1*A*). The MIRKO mice and the insulin receptor-loxP (IRlox) control mice were fed with either a normal chow diet (ND) or a high fat diet (HFD; 60% Kcal from fat) for 24 weeks to induce obesity, followed by analysis of DIO-induced metabolic disorders. Although IR deficiency in skeletal muscle did not affect weight gain, the MIRKO mice exhibited increased content in fat mass relative to the IRlox control mice (Figure 1*B* and 1*C*). As expected, the MIRKO mice developed severe insulin resistance and glucose intolerance, which is supported by results from glucose and insulin tolerance tests (Figure 1*D* and 1*E*). Moreover, the MIRKO mice also exhibited with hyperinsulinemia, hypertriglyceridemia, and elevated the levels of serum total ketone bodies (TKB) both under ND and HFD (Figure 1*F*-1*H*).

**Figure 1.**
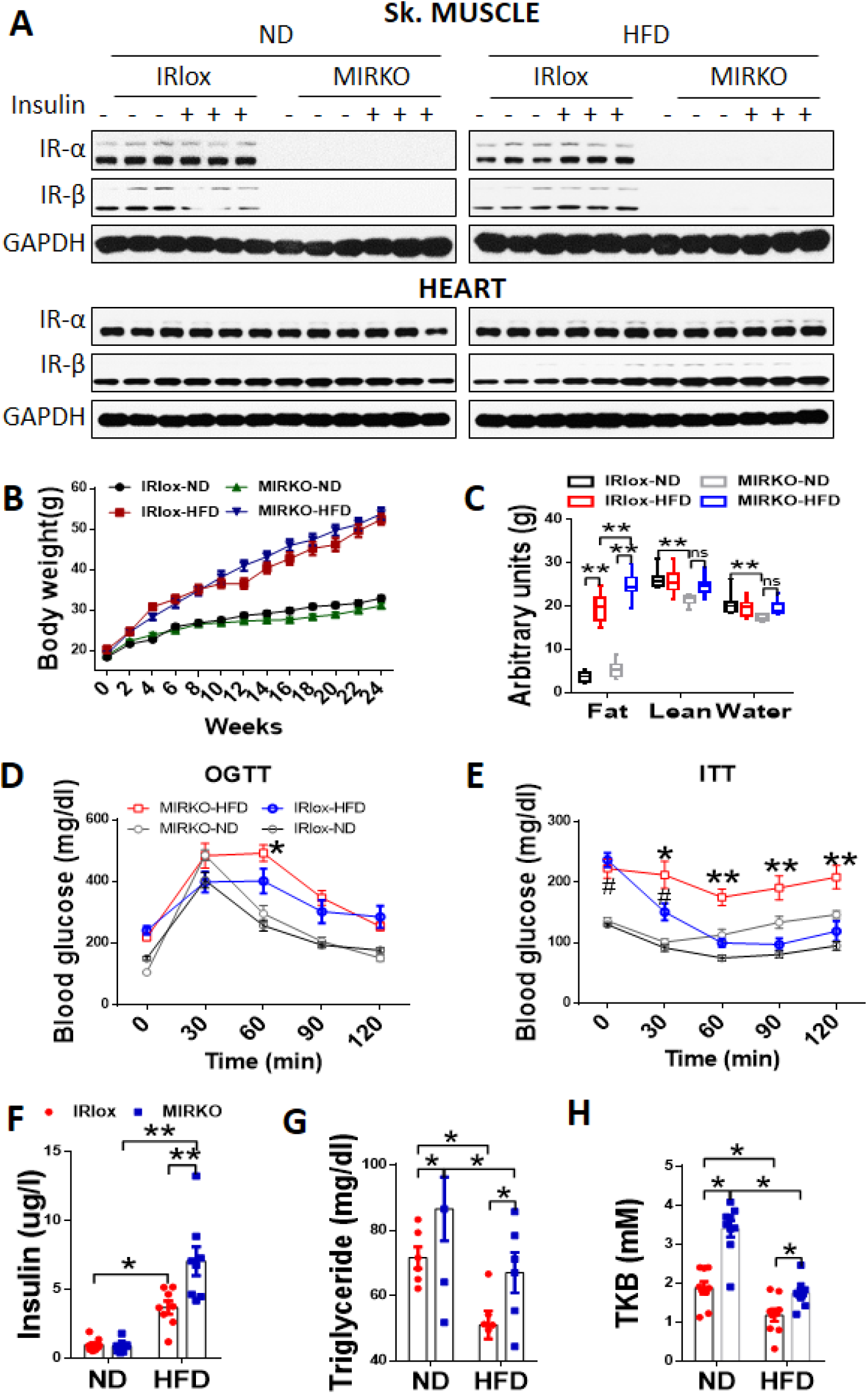
MIRKO aggravates DIO-induced insulin resistance, leading to hyperinsulinemia, glucose and insulin intolerance. (*A*) Evaluation of the insulin receptor knockout efficiency by immunoblot analysis of IR-α and IR-β, n=3 per group. (*B*) Body weight of MIRKO and IRlox control mice during ND and HFD feeding, n=10 per group. (*C*) Evaluation of whole-body composition using a quantitative magnetic resonance imaging system, n=10 per group. (*D* and *E*) Glucose and insulin tolerance tests in MIRKO and IRlox control mice either ND or HFD feeding, n=8-10. (*F*-*H*) Enzyme linked immunosorbent assay of the levels of insulin (*F*), triglycerides (*G*) and total ketone bodies (TKB) (*H*) in serum of MIRKO and IRlox control mice in response to DIO, n=8-10. GAPDH was used as internal control for protein loading. *, IRlox-HFD vs. MIRKO-HFD; ^#^, IRlox-HFD vs. IRlox-ND. Data are expressed as means ± SEM; ns, nonsignificant, *p<0.05, **p<0.01, ^#^p<0.05.

### Insulin resistance in skeletal muscle prevents DIO-induced LV dysfunction and adverse remodeling

We next determined the effect of skeletal muscle insulin resistance on DIO-induced left ventricle (LV) dysfunction and adverse remodeling. As demonstrated by a representative echocardiograph (Figure 2*A*), the onset of DIO caused LV contractile dysfunction, as shown by decreased LV fractional shortening (LVFS), LV ejection fraction (LVEF), LV posterior wall end-systole (LVPWs), and interventricular septal end-systole (IVSs) without significant effect on other parameters, including interventricular septal end-diastole (IVSd), left ventricular internal diameter end-diastole and end-systole (LVIDd and LVIDs), and LV posterior wall end-diastole (LVPWd) (Figure 2*B*-2*E*, Figure S1*A*-S1*D*). Remarkably, insulin resistance protected the MIRKO mice from development of the LV dysfunction, as supported by changes in LVFS, LVEF, LVPWs and IVSs relative to the IRlox control mice (Figure 2*A*-2*E*). Consistent with the findings, DIO also caused cardiac hypertrophy in IRLox control mice, which is evidenced by results from wheat germ agglutinin (WGA) staining of cardiac sections (Figure 2*F*, quantified in Figure 2*G*) and RT-PCR analysis of mRNA expression levels of hypertrophic biomarkers, including atrial natriuretic factor (ANF), brain natriuretic peptide (BNP) and β-cardiac myosin heavy chain (β-MHC) (Figure 2*H*). Again, IR deficiency in skeletal muscle not only mitigated these defects, but also prevented the onset of myocardial fibrosis, which is supported by results from Masson’s trichrome staining of heart samples, morphometric analysis of collagen volume fraction (CVF%) (Figure 2*I*-2*J*), and RT-PCR analysis of mRNA expression levels of both collagen I and collagen III (Figure 2*K*).

**Figure 2.**
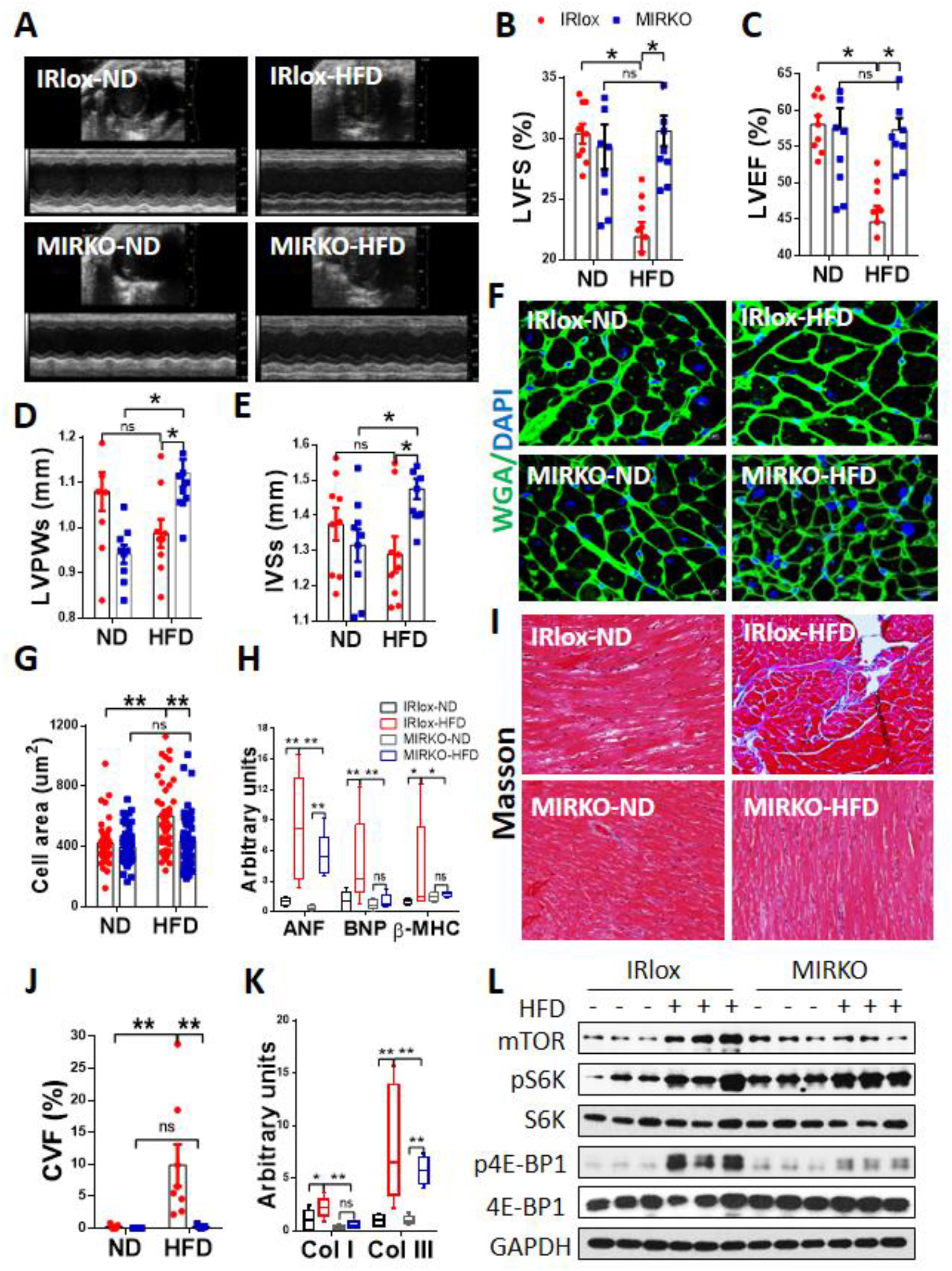
MIRKO mitigates DIO-induced LV dysfunction and adverse remodeling. Male skeletal muscle-specific insulin receptors knockout (MIRKO) and insulin receptor LoxP (IRlox) control mice were fed with normal diet (ND) or high fat diet (HFD) for 24 weeks. (*A*) Representative M-model echocardiographic images, n=8-10. (*B-E*) Summary data of echocardiographic parameters, including left ventricular fractional shortening (LVFS) (*B*), left ventricular ejection fraction (LVEF) (*C*), left ventricular posterior wall end-systole (LVPWs) (*D*) and interventricular septal end-systole (IVSs) (*E*), n=8-10. (*F* and *G*) Representative images of wheat germ agglutinin staining (WGA) (*F*), and quantification of myocardial area (*G*) of heart section from IRlox and MIRKO control mice in response to DIO, n=3 per group. (*H*) RT-qPCR analysis of the mRNA levels of myocardial hypertrophy biomarkers (ANF, BNP and β-MHC), n=6 per group. (*I* and *J*) Evaluation of the level of cardiac fibrosis by Masson’s trichrome staining (200x magnification) (*I*), and quantitative analysis of collagen volume fraction percent (CVF %) (*J*), n=3 per group. (*K*) RT-qPCR analysis of the mRNA levels of collagen I (col I) and III (col III), n=6 per group. (*L*) Immunoblot analysis of the phosphorylation levels of the mTOR signaling pathway in heart tissues, including S6K and 4E-BP1. GAPDH was used as internal control for mRNA and protein loading. Data are expressed as means ± SEM; ns, nonsignificant, * p<0.05, **p<0.01.

The mammalian target of rapamycin (mTOR) signaling pathway controls cell growth and size. Hyperactivation of mTORC1 is often implicated in the pathogenesis of cardiac hypertrophy (14). We next determined whether insulin resistance in skeletal muscle would attenuate hyperactivation of mTORC1 signaling in the heart of MIRKO mice in response to DIO. Consistent with cardiac hypertrophy, the mTORC1 signaling pathway in the heart was significantly activated by DIO in IRlox control mice, which is supported by both increased mTOR protein expression level and phosphorylation levels of its downstream targets, including S6K and 4E-BP1 (Figure 2*L*, quantified in Figure S1*E*). In further support of a protective role of skeletal muscle insulin resistance to the heart, MIRKO normalized the phosphorylation levels of both mTOR and 4E-BP1 without significant effect on S6K signaling in the heart.

### MIRKO selectively mitigates DIO-induced insulin resistance in heart, but not in other metabolic tissues

To gain insight into the molecular mechanisms by which skeletal muscle insulin resistance improved cardiac function, we next investigated the effect of MIRKO on insulin signaling in major metabolic tissues, including heart, skeletal muscle (Sk. muscle), fat, and liver. As expected, insulin stimulated phosphorylation of AKT and GSK3β were totally blunted in skeletal muscle when mice were fed with both ND and HFD in MIRKO mice, which is supported by decreased levels of p-AKT (T308), p-AKT (S473), and p-GSK3β (Figure 3*A*, quantified in Figure 3*B*). Strikingly, in spite of hyperinsulinemia, insulin sensitivity was significantly improved in the heart of MIRKO mice in response to DIO, as evidenced by increased phosphorylation levels of AKT-threonine-308 (T-308), a key phosphorylation site that mediates insulin sensitivity (Figure 3*A*, quantified in Figure 3*C*) (15). In contrast, hyperinsulinemia in MIRKO mice significantly impaired insulin sensitivity in other metabolic tissues, as suggested by decreased phosphorylation levels of AKT in adipose tissue and liver (p-AKT, T308 and S473) and GSK3β (Figure S2*A*, quantified in S2*B* and S2*C*). Together, these findings suggest that insulin resistance in skeletal muscle mediates a specific crosstalk with the heart, but not other metabolic tissues, in response to metabolic stress in DIO.

**Figure 3.**
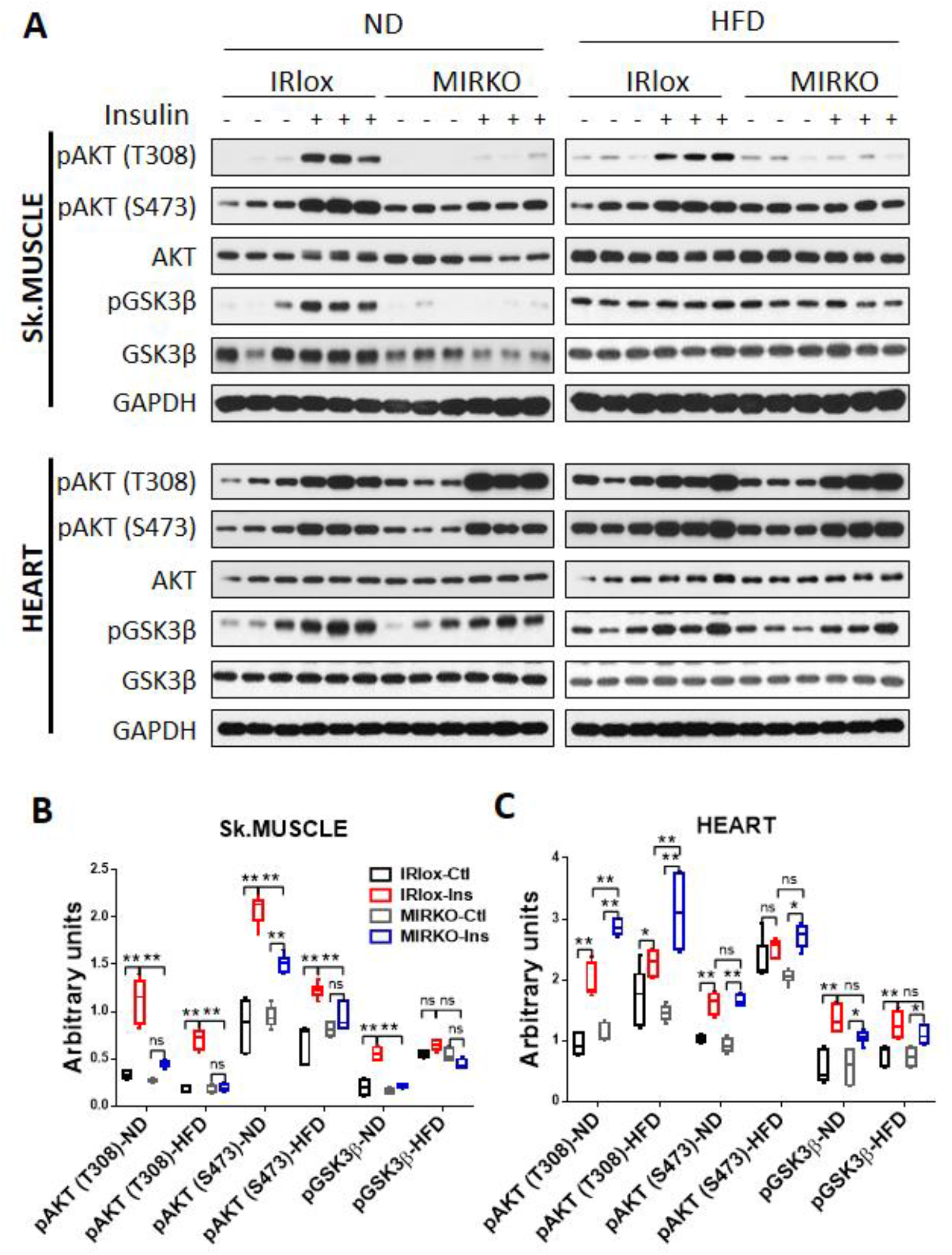
MIRKO selectively mitigates DIO-induced insulin resistance in heart. (*A*) Immunoblot analysis of AKT phosphorylation at Thr308 and Ser473, and GSK3β phosphorylation in skeletal muscle and heart tissues from MIRKO and IRlox control mice after i.p. injection of 1 U/kg insulin for 15 min, n=3 per group. (*B* and *C*) Quantitative analysis of the phosphorylation levels of AKT and GSK3β. GAPDH was used as internal control for protein loading. Data are expressed as means ± SEM ; ns, nonsignificant, *p<0.05, **p<0.01.

### MIRKO prevents DIO-induced inflammation and apoptosis in heart

The onset of DIO causes chronic inflammation (16, 17), which is implicated in adverse cardiac remodeling (18). To gain further insight into the molecular mechanisms underlying the protective effects, we next investigated whether skeletal muscle insulin resistance would prevent myocardial inflammation and apoptosis in MIRKO mice. Accordingly, we showed that the onset of DIO led to activation of multiple inflammatory pathways, including NLRP3, TXNIP, and cGAS-cGAMP-STING pathways, as evidenced by results from western blot analysis of key inflammatory biomarkers in the hearts of the IRlox control mice (Figure 4*A*, quantified in 4*B*). Our findings are corroborated by previous reports that NLRP3, an intracellular sensor for various “danger” signals, is activated by the onset of DIO (16, 17) and by adverse cardiac remodeling (19). TXNIP is an activator of NLRP3 and a key sensor of oxidative stress, and targeted deletion of TXNIP protects the myocardium from ischemia-reperfusion injury (20). Although the cGAS-cGAMP-STING pathway is primarily involved in antiviral responses, recent studies show that it can also be activated by cytosolic release of mitochondrial DNA (mtDNA) from metabolic stress, and therefore ablation of STING significantly attenuated chronic inflammation in DIO (17). Consistent with beneficial effects to the heart, the MIRKO mice are protected from myocardial inflammation induced by DIO, as evidenced by decreased protein expression levels of major inflammatory biomarkers (Figure 4*A*, quantified in 4*B*). Consistent with the findings, MIRKO also attenuated the mRNA expression levels of multiple pro-inflammatory cytokines, including TNF-α, NF-kB, IL-1β and IL-6, which were significantly elevated in the hearts of the IRlox control mice in response to DIO (Figure 4*C*). Consequently, the MIRKO mice were also protected from apoptosis of cardiomyocytes induced by DIO, which is supported by the normalized protein expression levels of major apoptotic biomarkers, including BAX, cleaved caspase 3 (C-cas3), cytochrome c (Cyto c), and Bcl-2, as well as by results from TUNEL staining of heart samples (Figure 4*A* and 4*E*, quantified in Figure 4*D* and 4*F*).

**Figure 4.**
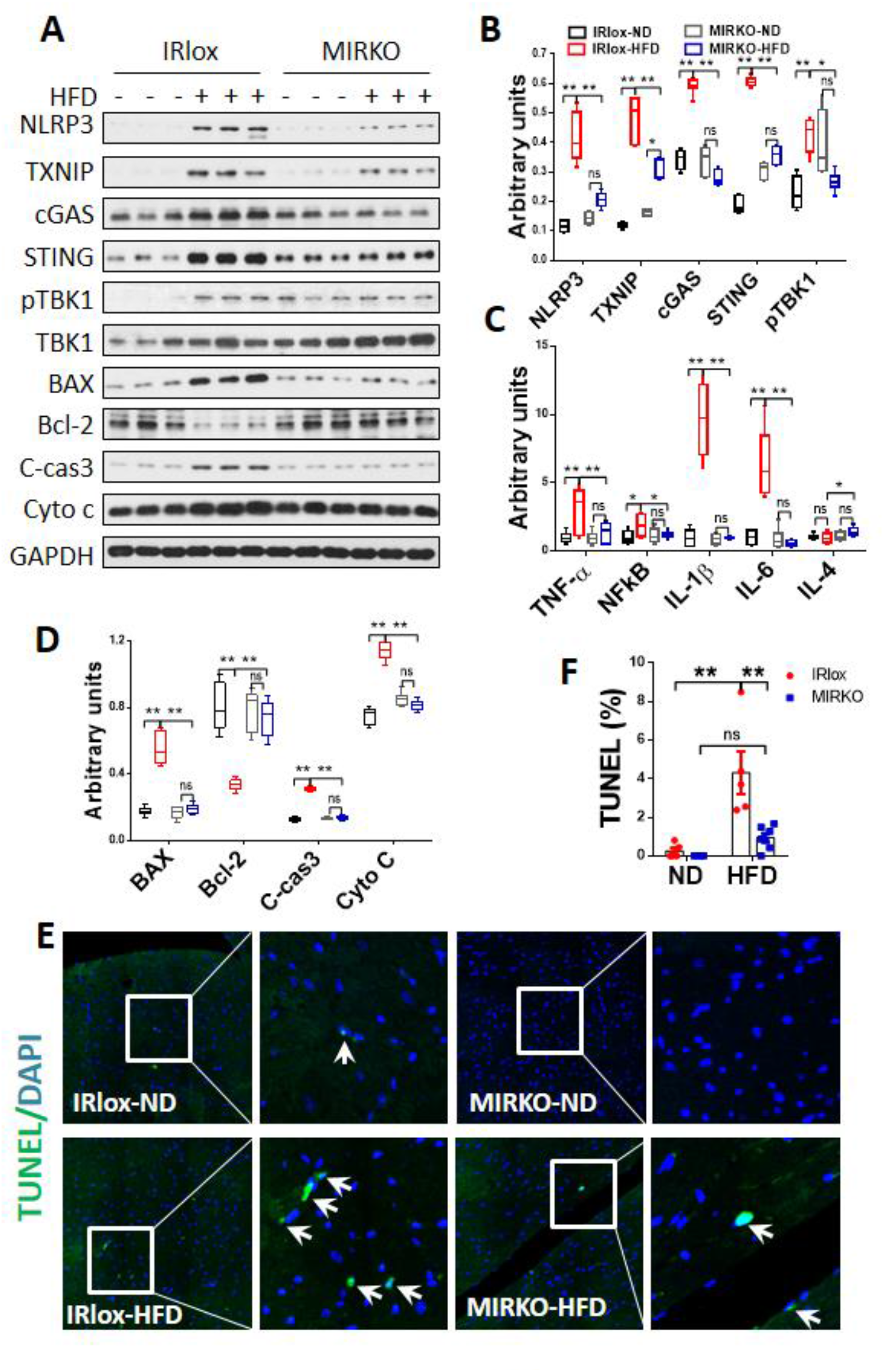
MIRKO prevents DIO-induced inflammation and apoptosis in heart. (*A*) Immunoblot analysis of inflammation and apoptosis associated biomarkers, including NLRP3, TXNIP, STING signal pathway (cGAS, STING, TBK1), and apoptosis (BAX, Bcl-2, C-cas3, Cyto c), n=3 per group. (*B*) Quantitative analysis of the protein expression of inflammation biomarkers. (*C*) RT-qPCR analysis of the mRNA levels of anti-inflammation (IL-4) and pro-inflammation (TNF1-α, NFKB, IL-1β, IL-6) cytokines, n=6 per group. (*D*) Quantitative analysis of the protein expression of apoptosis (BAX, Bcl-2, C-cas3, Cyto C) biomarkers. (*E* and *F*) Representative images (*E*) and quantitative analysis (*F*) of apoptosis in heart sections stained with TUNEL in the MIRKO and IRlox control mice in response to DIO. Arrows highlight apoptotic cells. n=3 per group. GAPDH was used as internal control for mRNA and protein loading. Data are expressed as means ± SEM; ns, nonsignificant, * p<0.05, **p<0.01.

### MIRKO mitigates DIO-induced myocardial lipid accumulation by promoting mitochondrial fatty acid oxidation

Myocardial insulin resistance causes mitochondrial dysfunction, which is implicated in the pathogenesis of myocardial hypertrophy (21, 22). As one of the highest energy demanding organs per gram of tissue weight in the body, the heart heavily depends on mitochondrial fatty acid oxidation (FAO) to provide high-density fuel. During adverse cardiac remodeling, cardiac metabolism is reprogrammed toward increased reliance on glycolysis, whereas FAO is downregulated (23). To further identify mechanisms underlying the beneficial effects of skeletal muscle insulin resistance to the heart, we next investigated whether skeletal muscle insulin resistance would prevent the metabolic reprogramming in the heart of the MIRKO mice. As shown by results from electron microscopic analysis of the heart samples, DIO caused significant accumulation of mitochondrial associated lipid droplets (LDs) in the heart of the IRlox control mice (Figure 5*A*, quantified in Figure 5*B*). Accumulation of mitochondria bound lipid droplets is associated with impaired fatty acid oxidation (24). Consistent with the notion, DIO also significantly increased intramyocardial triglyceride content, suggesting a defect in fatty acid oxidation (Figure 5*C*). In further support of the notion, the onset of DIO also caused depletion of hydroxyacyl-CoA dehydrogenase trifunctional multienzyme complex subunit-β (HADHB), which catalyzes the last three steps of mitochondrial β-oxidation of long chain fatty acids in heart and skeletal muscle tissues (Figure 5*D*, quantified in 5*E*-5*F*). In contrast, MIRKO not only prevented accumulation of myocardial mitochondria bound lipid droplets (Figure 5*A*, quantified in 5*B*), but also specifically restored HADHB protein expression, leading to a significant decrease in triglyceride content in the heart relative to other metabolic tissues, including skeletal muscle, liver, and fat (Figure 5*D* and Figure S3*A*, quantified in 5*E*-5*F* and S3*B*-S3*C*).

**Figure 5.**
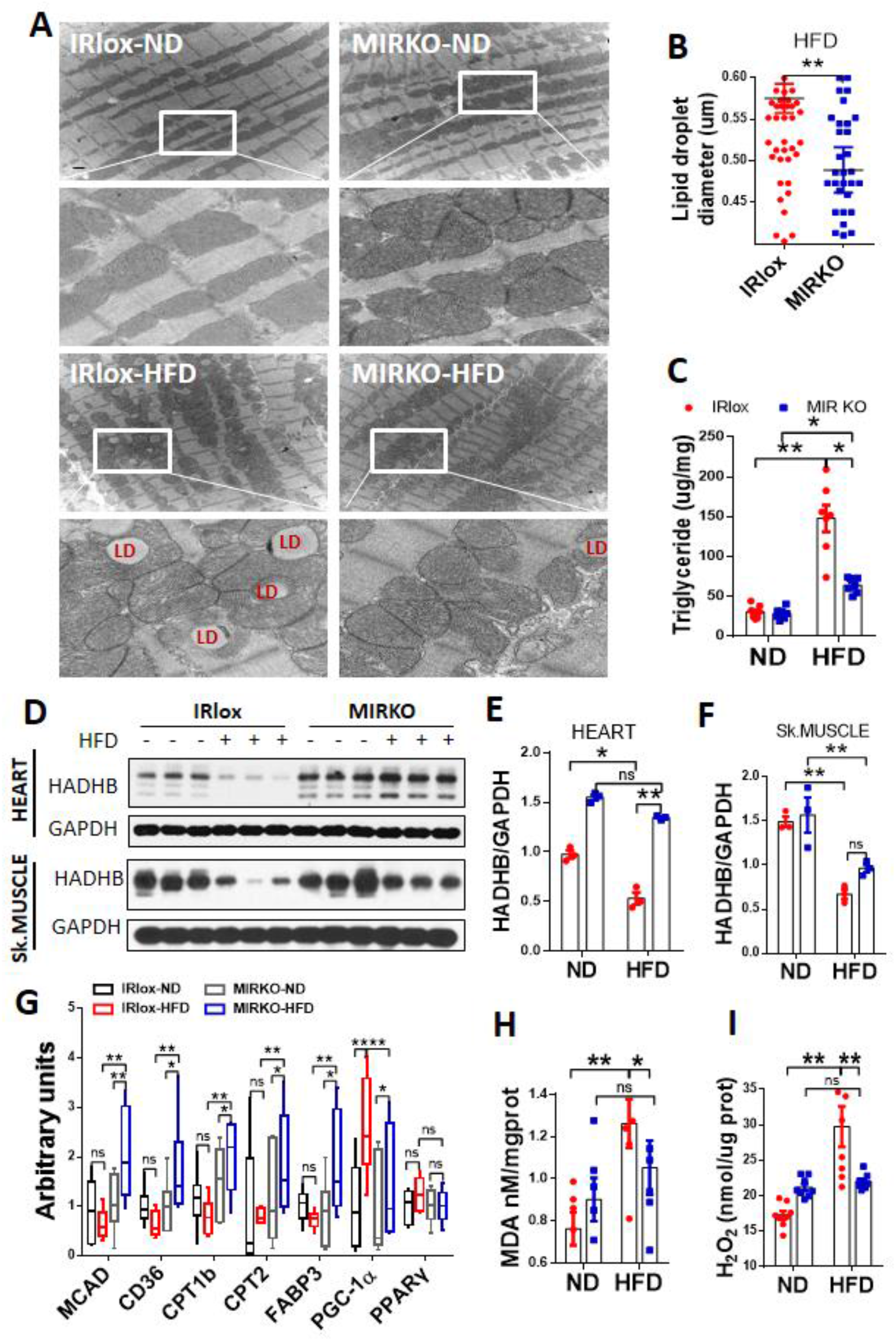
MIRKO mitigates DIO-induced myocardial lipid accumulation by promoting mitochondrial fatty acid oxidation. (*A*) Representative images of transmission electron microscopy (TEM) performed in heart tissues of MIRKO and IRlox control mice in response to DIO, Scale bars represent 1 μm, n=2 per group. (*B*) Quantitative analysis of the diameter of lipid droplets (n=59 lipid droplet) in the left ventricle. (*C*) Evaluation of the level of triglycerides in heart tissues from MIRKO and IRlox control mice in response to DIO, n=6 per group. (*D*) Immunoblot analysis of mitochondrial trifunctional protein β-subunit (HADHB) in heart and skeletal muscle tissues from MIRKO and IRlox control mice in response to DIO, n=3 per group. (*E* and *F*) Quantitative analysis of the protein expression of HADHB in heart (*E*) and skeletal muscle (*F*) tissues, n=3 per group. (*G*) RT-qPCR analysis of the mRNA levels of mitochondrial fatty acid oxidative biomarkers (MCAD, CD36, CPT1b, CPT2, FABP3, PGC-1α, PPARγ) in heart tissues from MIRKO and IRlox control mice in response to DIO, n=6 per group. (*H* and *I*) Analysis of lipid peroxidation using malondialdehyde (MDA) levels (*H*) and oxidative stress (*I*) in heart tissues, n=6 per group. GAPDH was used as internal control for mRNA and protein loading. Data are expressed as means ± SEM; ns, nonsignificant, * p<0.05, **p<0.01.

In further support of a protective role of increased FAO to the heart, insulin resistance in skeletal muscle also significantly up-regulated mRNA expression of major genes encoding FAO enzymes in the heart of MIRKO mice, including medium-chain acyl-Coenzyme A dehydrogenase (MCAD), fatty acid translocase (CD36), carnitine palmitoyltransferase 1b and 2 (CPT1b and CPT2), fatty acid binding protein 3 (FABP3), peroxisome proliferator activated receptor gamma co-activator 1α (PGC1α), and peroxisome proliferator activated receptor gamma (PPARγ) (Figure 5*G*). MCAD is required for the initial step of β-oxidation. CD36 and FABP regulate membrane uptake of fatty acids, whereas CPT1 and CPT2 catalyze mitochondrial transport of fatty acids, a key step involved in FAO (25). PGC-1α is a coactivator of the PPARγ, a key transcription factor of genes encoding lipid metabolic enzymes in the heart (26). Moreover, DIO also caused mitochondrial dysfunction in the heart of IRlox control mice, as evidenced by decreased mRNA expression levels of key regulators of mitochondrial biogenesis and oxidative phosphorylation, including transcription factor A (TFAM), pyruvate dehydrogenase kinase-isoenzyme 4 (PDK4), and NADH dehydrogenase (ubiquinone) 1 alpha subcomplex 9 (Ndufa9) without effect on estrogen related receptor alpha (ERRα) and ubiquinol-cytochrome c reductase core protein 1 (Uqcrc1) (Figure S3*F*). In contrast, MIRKO not only restored the mRNA levels of these mitochondrial regulators, but also increased mtDNA copy number and citrate synthase activity, a key biomarker for the presence of intact mitochondria in the heart of MIRKO mice in response to DIO (Figure S3*D*-S3*E*). Consistent with the findings, MIRKO also significantly attenuated oxidative stress, as suggested by decreased the levels of malondialdehyde (MDA) and hydrogen peroxide (H_2_O_2_) (Figure 5*H* and 5*I*). Together, our findings suggest that insulin resistance in skeletal muscle prevents DIO-induced myocardial dysfunction in part by promoting mitochondrial FAO, which is corroborated by previous reports that myocardial insulin resistance caused metabolic switch from FAO to glucose metabolism, leading to oxidative stress and mitochondrial dysfunction (27, 28).

### MIRKO dramatically alters mRNA level of major myokines in skeletal muscle

Then comes the question how insulin resistance in the MIRKO mice mediated specific crosstalk with the heart? As the largest organ of the body in non-obese individuals, skeletal muscle is now considered to be an active endocrine organ releasing a host of myokines which are part of a complex network that mediates communication between muscle and other organs, including liver, fat, and heart tissues. Although certain myokines are implicated in the communication with the heart, little is known about the effect of insulin resistance on their production. In search of myokines as a possible mediator of the crosstalk between skeletal muscle and the heart, we analyzed the effects of MIRKO on mRNA expression of several myokines which are implicated in cardiac protection, including myonectin and myostatin. Myonectin was reported to protect the heart from ischemia-reperfusion injury, whereas myostatin prevented the metabolic switch from FAO to glycolysis during heart failure (29, 30). We showed that the onset of DIO caused depletion of mRNA levels of myonectin and myostatin. Strikingly, insulin resistance mimicked the effect of DIO on myokine production, leading to a total depletion of mRNA expression levels of both myonectin and myostatin in the skeletal muscle of MIRKO mice under chow diet, revealing an unexpected role of insulin signaling in regulating myokines production (Figure 6*A*-6*B*). Consistent with up-regulated levels of inflammatory cytokines, MIRKO also significantly up-regulated mRNA expression levels of interleukin-6 (IL-6) without any major effect on other myokines, including insulin-like growth factor α (IGF1α), fibroblast growth factor 21 (FGF21), and follistatin-like 1 (FSTL1) in the skeletal muscle of MIRKO mice (Figure 6*C*-6*F*). However, MIRKO failed to restore mRNA expression of these myokines in MIRKO mice, suggesting that these myokines are not likely to convey the signal from skeletal muscle to the cardiomyocytes.

**Figure 6.**
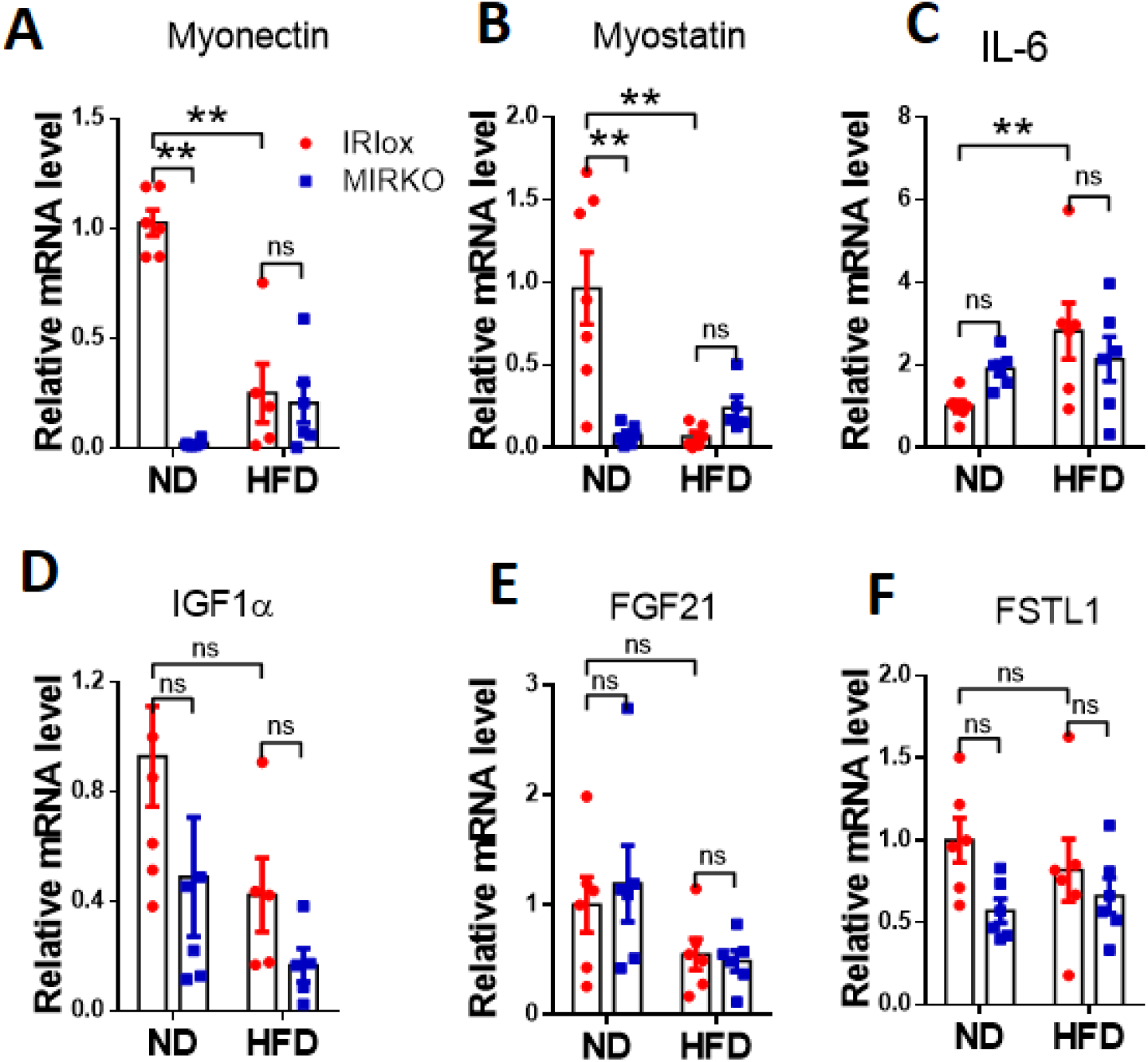
MIRKO dramatically alters mRNA level of major myokines in skeletal muscle. (*A*-*F*) RT-qPCR analysis of the mRNA expression of several myokines which are implicated in cardiac protection, including myonectin (*A*), myostatin (*B*), interleukin-6 (IL-6) (*C*), insulin growth factor α (IGF1α) (*D*), Fibroblast growth factor 21 (FGF21) (*E*), and follistatin-like 1 (FSTL1) (*F*) in the skeletal muscle, n=6 per group. GAPDH was used as internal control for mRNA loading. Data are expressed as means ± SEM; ns, nonsignificant, * p<0.05, **p<0.01.

## Discussion

Insulin resistance is implicated as the common cause of various chronic diseases, such as T2DM, diabetic complications, inflammation, and cardiovascular diseases. However, cumulative efforts in targeting insulin resistance for the treatment of these chronic conditions have not been met with great success to date. Consequently, the prevalence of heart failure remains very high to date, despite the success of many commonly used antihyperglycemic therapies to control hyperglycemia in T2DM in the last couple of decades (2). Although treatment of T2DM patients with thiazolidines, one of the most effective insulin sensitizing drugs, significantly improves glycemic control, these drugs are no longer widely prescribed in the clinic due to their unwanted cardiac side effects (7–10). Furthermore, it can be argued that if insulin resistance is truly a deleterious consequence of obesity, why has evolution not eliminated it? In this study, we addressed this dilemma by demonstrating that insulin resistance is protective to the heart in response to metabolic stress associated with DIO, which is supported by multiple lines of evidence. Despite exacerbating obesity and its related metabolic syndromes, the MIRKO mice were protected from development of LV hypertrophy, fibrosis, and apoptosis in response to DIO. These findings are further underscored by underlying changes in signal transduction pathways that mediates these pathogeneses, including the hyperactivation of mTORC1, NLRP3, STING, and apoptotic pathways in the heart.

Obesity causes myocardial insulin resistance, which is implicated in the pathogenesis of heart failure, as evidenced by the phenotype of mice with a cardiomyocyte-selective insulin receptor knockout (CIRKO). Therefore, it is commonly accepted that insulin resistance contributes to the development of cardiac hypertrophy and dysfunction in obese patients. However, this simplistic view is challenged by our findings that skeletal muscle insulin resistance is protective to the heart in response to DIO. Although hyperinsulinemia did exacerbate insulin resistance in liver and adipose tissues, MIRKO mice were protected from cardiac insulin resistance and mitochondrial dysfunction in response to DIO. We further demonstrated that the MIRKO mice also exhibited with a metabolic switch from glucose metabolism to FAO in the heart. This change was accompanied by upregulated expression of mitochondrial enzymes and proteins involved in FAO and oxidative phosphorylation, leading to significant attenuation of oxidative stress, lipid peroxidation, and accumulation of mitochondrial bound lipid droplets in the heart. These metabolic changes offer a possible explanation why the MIRKO mice were protected from DIO-induced myocardial hypertrophy and dysfunction. Consistent with this notion, the cardiac metabolism is reprogrammed toward increased reliance on glycolysis, whereas FAO is downregulated in response to adverse myocardial remodeling and heart failure (23). Our findings are also corroborated by the phenotypes of CIRKO mice which exhibit myocardial insulin resistance, impaired FAO, and upregulated glycolysis in the heart (32).

Despite intensive research in the field, the potential benefit of insulin resistance, if any, in the survival of mammals remains elusive. Our findings revealed for the first time that insulin resistance in skeletal muscle mediates a specific crosstalk with the heart to prevent cardiac dysfunction under metabolic stress. This notion is indirectly supported by the phenotypes of various mouse models of targeted deletion of insulin receptor or glucose transporter-4 (GLUT-4) which plays a critical role in insulin receptor-mediated glucose uptake in all metabolic tissues. In contrast to MIRKO mice, mice with both skeletal muscle and cardiac muscle knockout of the GLUT4 gene (muscle-G4KO) developed glucose intolerance and insulin resistance as early as 8 weeks of age (31). Additionally, a subset of muscle-G4KO mice developed frank diabetes with severe insulin resistance. Moreover, in contrast to cardiac hypertrophy in muscle-G4KO mice, mice with cardiac specific knockout of GLUT-4 (cardiac-G4KO) exhibit normal contractile function and glucose homeostasis, suggesting a crosstalk between skeletal muscle and the heart in regulating glucose homeostasis as well (32). Strikingly, in contrast to MIRKO mice, neither the cardiac-G4KO nor the muscle-G4KO mice developed hyperinsulinemia and dyslipidemia, implicating an important role of metabolic syndrome in regulating cardiac function (3, 32). Intriguingly, this notion is also corroborated by clinical observations that overweight and obese patients with CVD have better prognoses compared with leaner patients with the same cardiovascular diagnoses (4–6). Since most obese patients have metabolic syndromes, the findings further contradict the commonly held belief that insulin resistance exacerbates heart failure.

Together, our findings revealed a highly unexpected role of insulin resistance in skeletal muscle as a double-edged sword. On one hand, it protected the heart from myocardial dysfunction in response to metabolic stress associated with DIO, on the other it exerted a deleterious effect on glucose homeostasis by exacerbating DIO-induced insulin resistance in other metabolic tissues, including fat and liver which could lead to the development of microvascular complications. Although molecular mechanisms underlying the crosstalk remains fully elucidated, our studies have provided the first glimpse into this complicated issue, calling for reevaluation of ongoing efforts in targeting insulin resistance for the treatment for metabolic diseases. If the mechanism(s) responsible for cardiac improvement can be separated from the negative impact on glucose tolerance, this would have enormous potential implications for the treatment of heart failure and other CVD. However, how the development of skeletal muscle insulin resistance transmits its message to the heart remains unclear. Likewise, why hyperinsulinemia failed to cause insulin resistance in the heart of MIRKO mice remains a fascinating question that deserves further investigation in the future.

## Materials and Methods

### Reagents

Antibodies used in the present studies include polyclonal antibodies to insulin receptor α (IR-α) and β (IR-β), phospho-AKT (pAKT, Thr308), phospho-AKT (pAKT, Ser473), AKT, Phospho-GSK3β (pGSK3β, Ser9), GSK3β, mTOR, phospho-S6K (pS6K, Thr389), S6K, phospho-4EBP1 (p4EBP, Thr37/46) and 4EBP1, NLRP3, TXNIP, cGAS, STING, phospho-TBK1 (pTBK1), TBK1, BAX, Bcl-2, Cleaved Caspase3 (C-cas3) and Cytochrome c (Cyto c) all of which were purchased from Cell Signaling Technology (Danvers, MA). Anti-HADHB antibody was from Abcam. Mouse anti-GAPDH antibody were from Santa Cruz Biotechnology (Santa Cruz, CA). Bicinchoninic acid (BCA) protein Assay Kit was from Pierce (Rockford, IL, USA).

### Animal care

The skeletal muscle-specific insulin receptor knockout mice (MIRKO) were generated by crossing insulin receptor loxP mice (IRlox) (Jackson Laboratory; stock No. 006955) with skeletal muscle-Cre mice (Jackson Laboratory; stock No. 024713). Genotyping was performed by PCR using genomic DNA isolated from the tip of the tail of 3-to 4-week-old mice. Six-week-old male MIRKO and IRlox control mice were fed with a normal chow diet (ND) or a high fat diet (HFD; 60% Kcal from fat, Research Diets) for 24 weeks. All animals were maintained in an environmentally controlled facility with a diurnal light cycle and free access to water and a standard rodent chow. All experiments involving animals were performed in compliance with approved institutional animal care and use protocols according to NIH guidelines (NIH publication no.86-23 [1985]).

### Body Composition Analyses

Mice body weight was measured once every two weeks. MIRKO and IRlox control mice were fed with ND or HFD for 24 weeks. Measurements of fat, lean, and water masses was determined using EchoMRI™.

### Insulin and Glucose Tolerance Test

In brief, insulin tolerance test (ITT) was performed in 6-hrs fasting mice. Mice were subjected to insulin (1.0 U/kg body weight; Novolin, Novo Nordisk) by intraperitoneal injection (i.p.). Oral glucose tolerance test (OGTT) was performed on mice fasted overnight. Mice were gavaged with glucose (2.5 g/kg body weight) and blood glucose was determined at 0, 30, 60, and 120 mins with a One Touch Ultra 2 glucometer (Lifescan, Milpitas, California).

### Insulin, Total Ketone Bodies (TKB), and Triglyceride Analyses

Insulin in serum was analyzed by Mouse Insulin ELISA kit (Mercodia, 10-1247-01) according to manufacturer’s instruction. Total ketone bodies (TKB) in serum was analyzed by measuring 3-hydroxybutyric acid (BOH) and acetoacetic acid (AcAc) using Ketone body assay kit (Sigma, No. MAK134) according to manufacturer’s instruction. Triglyceride in serum was analyzed using Triglyceride quantification colorimetric/fluorometric kit (Sigma, No. MAK266) according to manufacturer’s instruction.

### Histological Analyses

Three mice from each group were anesthetized with isoflurane. Heart samples were isolated and fixed with 4% paraformaldehyde for 48 hours. Fixed hearts were dehydrated and embedded in paraffin, and 5 μm sections were cut with a Leica RM-2162 (Leica Microsystems, Bensheim, Germany). Hematoxylin and Eosin (H&E) and Masson’s trichrome staining were performed according to manufacturer’s instruction. The collagen volume fraction (CVF) was analyzed by Image J. Wheat germ agglutinin (WGA, Thermo Fish Scientific) staining was performed by using Alexa Fluor 488 conjugate dye.

### Echocardiography Assay

Echocardiographic analysis was performed after 24 weeks of HFD feeding. M-mode short axis and B-mode long axis images of the left ventricle were analyzed to measure the following parameters, including left ventricular ejection fraction (LVEF), left ventricular fractional shortening (LVFS), left ventricular internal diameter end-diastole and end-systole (LVIDd and LVIDs, mm), interventricular septal end-diastole and end-systole (IVSd and IVSs, mm), left ventricular posterior wall end-diastole and end-systole (LVPWd and LVPWs, mm).

### TUNEL staining and transmission electron microscopy (TEM) Assay

TUNEL staining was carried out by using an ApopTag® Fluorescein in Situ Apoptosis Detection Kit (S7110, Sigma) according to the manufacturer’s instruction. TUNEL positive cell percentage was analyzed using Image J.

The mitochondrial ultrastructure in mouse cardiomyocytes was evaluated using transmission electron microscopy (TEM). Heart samples from the same site of the left ventricle were fixed in 5% glutaraldehyde and 4% paraformaldehyde in 0.1 M sodium cacodylate buffer (pH 7.4) with 0.05% CaCl_2_ for 24 hours. After washing in 0.1 M sodium cacodylate buffer, tissues were post fixed overnight in 1% OsO4 and 0.1 M cacodylate buffer, dehydrated, and embedded in EMbed-812 resin. The sections were stained with 2% uranyl acetate, followed by 0.4% lead citrate, and viewed with a JEOL JEM-2200FS 200KV electron microscope (Electron Microscopy Sciences in University of Texas Health Science Center at San Antonio). Lipid droplet diameter and sarcomere length was analyzed by Image J.

### Mitochondrial DNA Copy Number Assay

Mitochondrial DNA (mtDNA) copy number was determined using a multiplexed real time PCR (RT-PCR) assay as previously described. Briefly, total DNA was extracted from the left ventricle using DNeasy Blood & Tissue Kit (QIAGEN, No. 69504) according to manufacturer’s instruction. RT-PCR analysis was carried out using SYBR Green Master Mix (Thermo Fisher, USA). Mitochondrial-specific (ND1) and Cyclophilin A control were determined in triplicated of each sample.

### ROS production, Lipid Peroxidation and Citrate Synthase Assay

The ROS production from isolated mitochondria from left ventricular homogenates was indirectly detected by quantitatively measuring hydrogen peroxide (H_2_O_2_) as previously described. Lipid peroxidation production in heart tissues was analyzed by measuring changes in malondialdehyde (MDA) using a TBARS kit (Cayman Chemical Company, Cat No.10009055) according to the manufacturer’s instruction. Citrate Synthase (CS) activity in heart tissue homogenate was measured by analyzing the production of SH-CoA by monitoring the absorbance of Citrate Synthase Developing Reagent using a Citrate Synthase Activity Assay kit (Cayman Chemical Company, No. 701040).

### Quantitative RT-PCR analysis

In brief, total RNA of a subset of heart tissues was extracted using TRIzol following the manufacturer instructions (Thermo Fisher, USA). 2 μg of total RNA was used for cDNA synthesis with Superscript II Reverse Transcriptase (Invitrogen). RT-qPCR analysis was carried out using SYBR Green Master Mix (Thermo Fisher, USA). The sequence of the primers used to detect these genes are listed in Supplemental Table 1.

### Statistical Analysis

Statistical comparisons were analyzed using two-tailed non-paired t-tests to evaluate the results by GraphPad Prism software (version 6.0). The values were considered statistically significant at p values of *p < 0.05, ** p < 0.01. Data are expressed as means ± SEM.

## Acknowledgements

We would like to thank Drs. Ralph DeFranzo and Nicolas Musi for critically reading this manuscript and for providing us with valuable feedback and suggestions on the experimental design. The authors acknowledge funding support from the National Institutes of Health (2R01DK076685-06A1, Y.S.), American Diabetes Association (1-14-BS-185, Y.S.), and a grant from the Barth Syndrome Foundation (to Y.S.).

## Author Contributions

D.J., J.Z., X.L. performed the experiments and analyzed the data; Y.S. designed the experiments and wrote the manuscript; J.N. and J.P.A. provided feedback and guidance on experimental designs. Z.T. provided mentorship to D.J. All authors reviewed results and commented on the manuscript.

---

**Figure S1.**
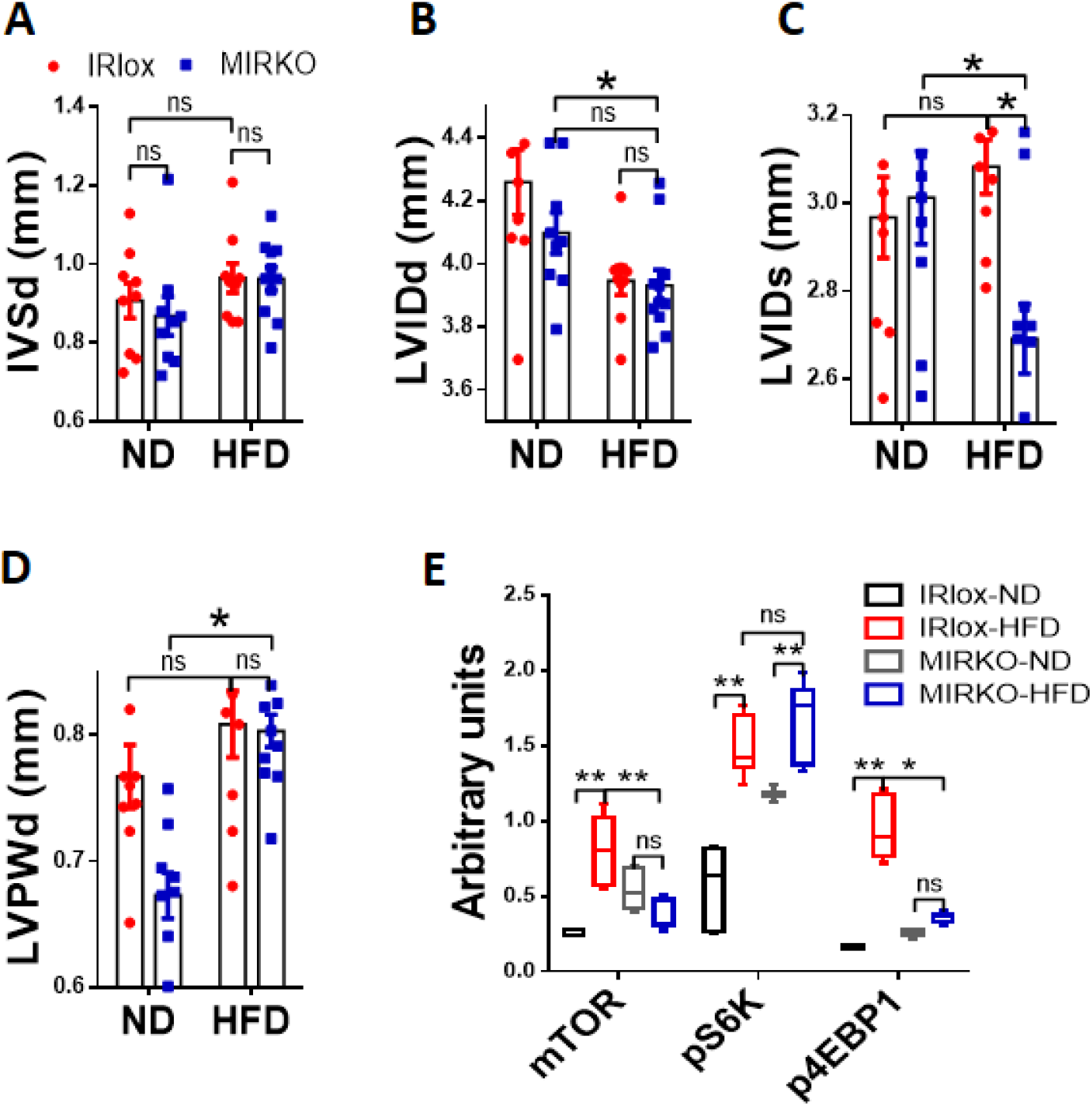
MIRKO mitigates DIO-induced LV dysfunction and adverse remodeling. (*A*-*D*) Summary data of echocardiographic parameters, including interventricular septal end-diastole (IVSd) (*A*), left ventricular internal diameter end-diastole and end-systole (LVIDd and LVIDs) (*B* and *C*), and left ventricular posterior wall end-diastole (LVPWd) (*D*). n=10 per group. (*E*) Quantitative analysis of the phosphorylation levels of the mTOR signaling pathway in heart tissues. n=3 per group. Data are expressed as means ± SEM; ns, nonsignificant, *p<0.05, **p<0.01.

**Figure S2.**
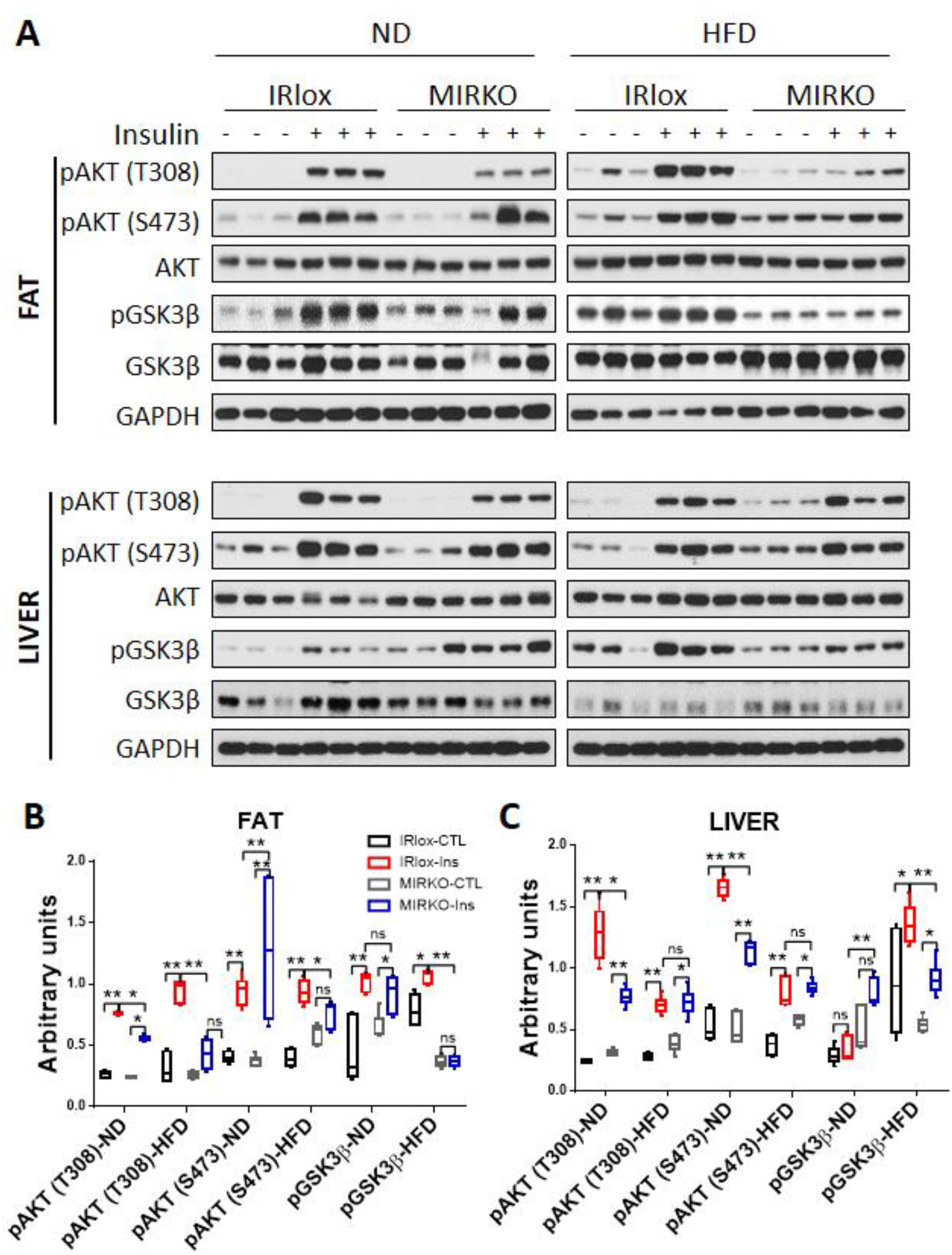
MIRKO aggravates DIO-induced insulin resistance in fat and liver. (*A*) Immunoblot analysis of AKT phosphorylation at Thr308 and Ser473 and GSK3β phosphorylation in fat and liver tissues from MIRKO and IRlox mice after i.p. injection of 1 U/kg insulin for 15 min, n=3 per group. (*B* and *C*) Quantitative analysis of the phosphorylation levels of AKT and GSK3β. GAPDH was used as internal control for protein loading. Data are expressed as means ± SEM; ns, nonsignificant, *p<0.05, **p<0.01.

**Figure S3.**
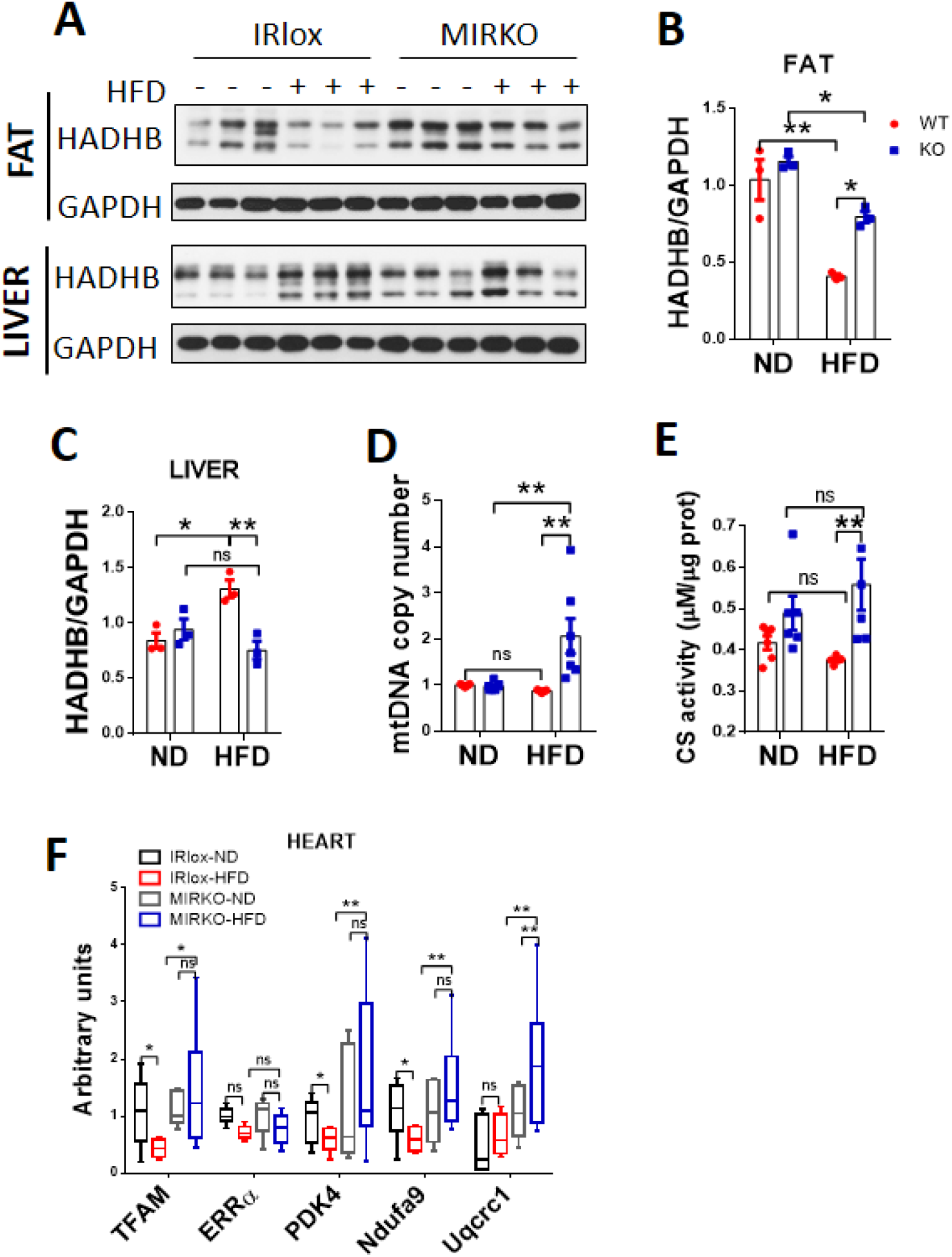
MIRKO promotes mitochondrial biogenesis and respiration in the heart. (*A*) Immunoblot analysis of mitochondrial trifunctional protein β-subunit (HADHB) in fat and liver tissues from MIRKO and IRlox control mice in response to DIO, n=3 per group. (*B*-*C*) Quantitative analysis of the protein expression of HADHB in fat (*B*) and liver (*C*) tissues. (*D*) RT-qPCR analysis of mtDNA copy number in heart tissues from MIRKO and IRlox control mice in response to DIO, n=6 per group. (*E*) Evaluation of the level of citrate synthase activity (CS) in heart tissues, n=6 per group. (*F*) RT-qPCR analysis of mitochondrial metabolism biomarkers (TFAM, ERRa, PDK4, Ndufa9, Uqcrc1) in heart tissues from MIRKO and IRlox control mice in response to DIO, n=6 per group. GAPDH was used as internal control for mRNA and protein loading. Data are expressed as means ± SEM; ns, nonsignificant, * p<0.05, **p<0.01.

**Supplemental Table 1.**
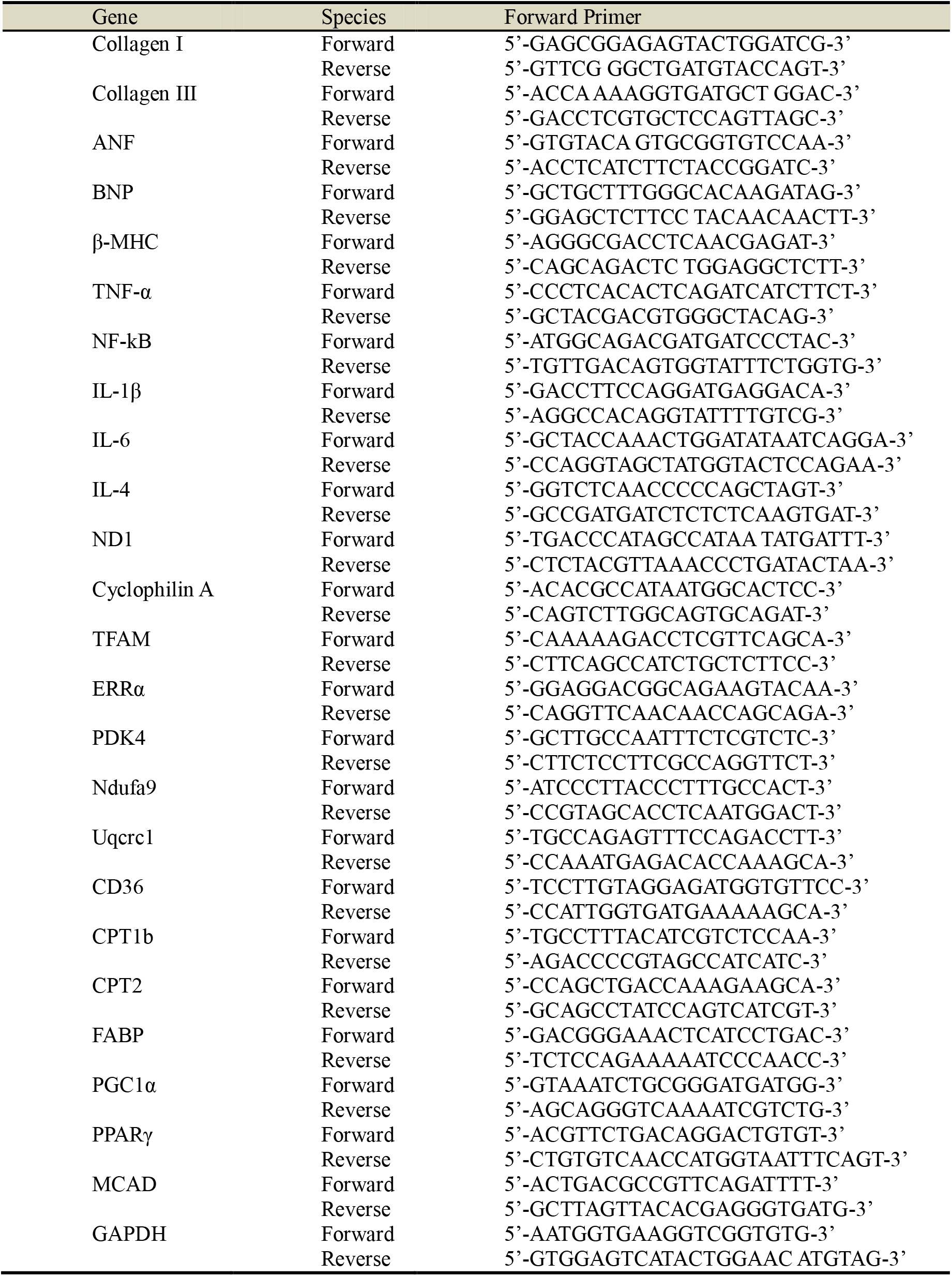
List of utilized primers for qRT-PCR

